# The exopolysaccharide Poly-N-Acetyl-Glucosamine (PNAG) coats *Klebsiella pneumoniae in vivo*

**DOI:** 10.1101/2024.09.23.614408

**Authors:** Jonathan Bradshaw, Julia Sanchez-Garrido, Rita Berkachy, Jaie Rattle, Connor Preston, Mariagrazia Pizza, Immaculada Margarit Ros, Maria Rosaria Romano, Joshua L.C. Wong, Gad Frankel

## Abstract

The conserved bacterial polysaccharide Poly-N-Acetyl-Glucosamine (PNAG) is a potential broad-spectrum vaccine candidate. While the immunogenicity of PNAG-based vaccine candidates has been established, characterisation of PNAG production across clinically relevant bacteria remains largely unknown. In particular, PNAG production in the Gram-negative pathogen *Klebsiella pneumoniae* (KP) is not well understood. Here, we demonstrate that PNAG production is prevalent in clinical KP isolates, where it is secreted as extracellular networks during adherent growth conditions. However, during severe KP pulmonary infection, KP PNAG production undergoes a switch to a cell-associated phenotype, coating the bacterial cell surface. By screening a panel of isogenic KP mutants in prominent cell surface components (**Δ***wcaJ*, **Δ***rmpADC*, **Δ***rfb*, **Δ***ompA* and **Δ***ompk36*), we identified KP capsular polysaccharide as a key determinant underpinning the phenotype. Deleting genes involved in capsule synthesis (**Δ***wcaJ*) and regulation (**Δ***rmpADC*) resulted in cell-associated PNAG during adherent growth and infection of alveolar epithelial cells *in vitro*. Taken together, we describe a novel interaction between KP surface polysaccharides and detect for the first time, cell-associated PNAG in KP during lung infection, highlighting PNAG as an attractive KP vaccine antigen.

**Author summary:** The Gram-negative pathogen *Klebsiella pneumoniae* (KP) is a leading cause of hospital-associated lung and bloodstream infections worldwide. As KP exhibits resistance to most frontline antibiotics, there is a growing demand for immune-based strategies to treat KP infections. Poly-N-Acetyl-Glucosamine (PNAG) is a surface sugar produced by most clinically relevant bacteria, including KP. However, relatively little is known about PNAG production in KP. Therefore, we set out to characterise PNAG production in KP during in vitro growth and following lung infection in a pulmonary mouse model. During *in vitro* growth, KP produces extracellular PNAG networks. In contrast, during an *in vivo* severe lung infection, PNAG is found cell-associated, coating the bacterial surface. We propose that the visible change in KP PNAG between *in vitro* and *in vivo* environments is due to crosstalk with capsule, another polysaccharide on the KP surface. Together, this supports PNAG as an attractive KP antigen.

## Introduction

Bacterial surface polysaccharides often contribute to virulence and represent attractive therapeutic targets. Poly-N-Acetyl-Glucosamine (PNAG) is a high molecular weight bacterial polysaccharide consisting of repeat units of N-Acetyl-Glucosamine (GlcNAc) joined via a β(1-6) glycosidic linkage [1,2]. Unlike other bacterial surface molecules, PNAG is conserved across bacteria, commensals and pathogens alike, and has been described in most members of World Health Organisation (WHO) critical and high priority pathogens [3–6]. As a result, PNAG is considered a broad-spectrum vaccine candidate and, over recent years, the immunogenicity of PNAG-based vaccine candidates has been investigated [7–9]. This has culminated in a recent study reporting that vaccination with PNAG glycoconjugates protects against lethal *Staphylococcus aureus* (SA) challenge in an *in vivo* model of peritonitis [10]. In SA, PNAG production is well characterised; however, PNAG production across most other bacterial pathogens remains relatively unexplored.

PNAG production in the encapsulated Gram-negative pathogen *Klebsiella pneumoniae* (KP) has not been well characterised. KP is a common constituent of the human microbiome, where it colonises the mucosal surfaces of the oropharynx and gastrointestinal (GI) tract [11]. Classical KP (cKP) strains cause infections in immunocompromised patients, commonly bacterial pneumonia and bloodstream infections, and are associated with broad antimicrobial resistance [12]. In contrast, hypervirulent KP (hvKP) strains can cause infections in healthy individuals and are associated with overproduction of the polysaccharide capsule and carriage of large virulence plasmids [13,14]. Infections caused by KP strains resistant to carbapenems and third generation cephalosporins are difficult-to-treat and have been assigned critical priority status by the WHO [3]. Therefore, there is a growing need for immune based therapies to prevent and treat KP infections.

In Gram-negative bacteria like KP, the genes required for PNAG synthesis are encoded on the *pgaABCD* locus [15]. During PNAG synthesis, an inner membrane complex forms between the glycosyltransferase PgaC and PgaD, a small protein with unknown function. The PgaCD complex polymerises UDP-GlcNAc to produce a fully acetylated PNAG polymer which extends into the periplasm [16,17]. In the periplasm, the deacetylase PgaB removes approximately 10% of the acetyl residues from the polymer to facilitate export through the outer membrane porin PgaA [18]. While the *pgaABCD* locus is prevalent across KP genomes, PNAG production has not been assessed across different KP strains [8]. Furthermore, while immunofluorescence staining has been successful in visualising PNAG production on many bacteria, attempts to visualise PNAG production on KP have yielded inconclusive results [4,6].

After being identified as a component of *Staphylococcal* biofilms, PNAG was found to contribute to virulence in diverse bacterial pathogens, including KP. Deletion of *pgaC* from a hvKP strain resulted in reduced colonisation of the gastrointestinal (GI) tract following oral inoculation and reduced lethality following peritoneal challenge [19]. Furthermore, prophylactic treatment with the PNAG-specific monoclonal antibody (mAb) F598 or polyclonal sera raised to PNAG antigen protected mice against intraperitoneal challenge with KP [6]. This is consistent with KP producing PNAG during infection of the peritoneum, and potentially being an attractive prophylactic and therapeutic target in KP peritonitis. However, the contribution of PNAG to KP virulence at other infection sites has not been explored. This includes during hospital-acquired pneumonia where KP is a leading cause [20].

In contrast to PNAG, other KP surface polysaccharides, capsule and lipopolysaccharide (LPS) O-antigen, have been extensively studied and now represent attractive vaccine and therapeutic targets for immunotherapy [21,22]. Therefore, we aimed to characterise KP PNAG production *in vitro* and during severe pulmonary infection in mice. We demonstrate that KP produces extracellular PNAG networks under adherent growth conditions *in vitro*, a phenotype conserved across diverse KP strains. However, we were unable to detect PNAG on strains belonging to the multilocus sequence type (MLST) ST258, a high-risk clonal group. We show that during severe pulmonary infection, KP undergoes a switch from extracellular network to cell-associated PNAG production. While immune therapies targeting extracellular networks of PNAG could act to perturb KP biofilm formation, cell-associated PNAG represents an attractive target to induce opsonic killing of KP. By deleting genes encoding capsule synthesis and regulatory proteins, we recapitulate this cell-associated PNAG phenotype during *in vitro* adherent growth conditions, and infection of alveolar epithelial cells. Collectively, this reveals a novel interaction between KP capsule and PNAG on the bacterial cell surface during severe pulmonary infection, and highlights PNAG as an attractive therapeutic target to treat KP pulmonary infection.

## Results

### KP produces extracellular PNAG networks during adherent growth conditions

While the *pga* operon has been identified in KP, little is known about PNAG production in this species. Therefore, we set out to visualise and characterise PNAG production across KP strains. We first aimed to visualise PNAG production in KP ICC8001 (our KP wild-type (WT)), a derivative of the prototype hvKP strain ATCC43816, which encodes an intact *pgaABCD* locus. We also generated an open reading frame deletion mutant in the *pgaC* gene (**Δ***pgaC*) as a negative control [15]. We assessed PNAG production by immunofluorescence staining of liquid cultures using the anti-PNAG mAb F598. However, we were unable to detect bacterial-associated or secreted PNAG under these growth conditions (Figure 1A).

**Figure 1.**
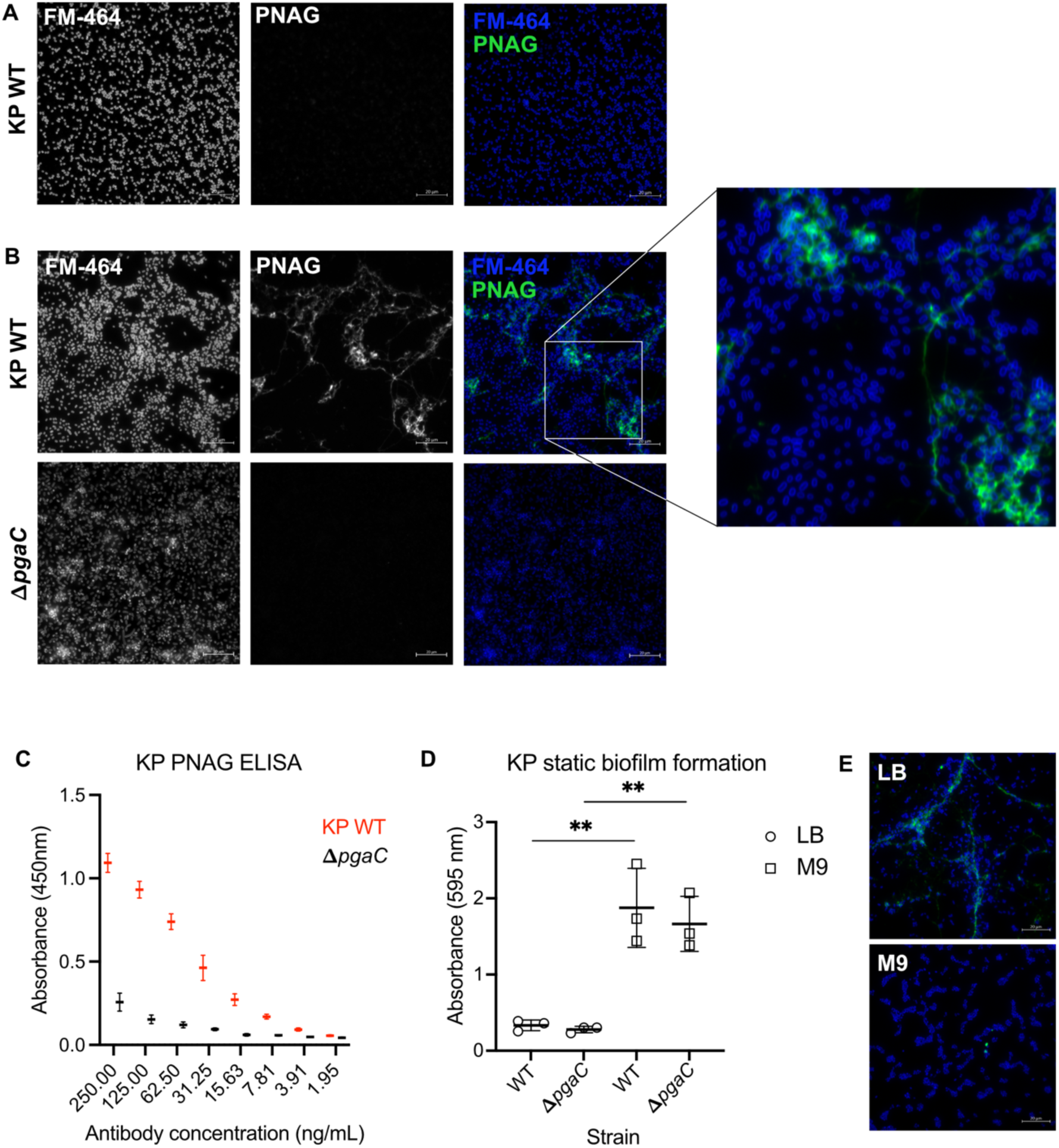
KP produces extracellular networks of PNAG during adherent growth conditions. **(A)** Visualisation of PNAG production in KP WT during growth in liquid media under agitation. KP WT was probed with mAb F598 to detect PNAG and the membrane dye FM4-64 to visualise bacteria (scale bar = 20µm). **(B)** Visualisation of PNAG production on KP WT and **Δ***pgaC* grown adherently on glass slides. A zoomed-in region of extracellular PNAG network formation is shown. **(C)** Quantification of PNAG production on adherent KP WT and **Δ***pgaC* following adherent growth. Data is presented as mean ±sd across 3 biological replicates. **(D)** Quantification of KP WT and **Δ***pgaC* static biofilm formation following static growth in nutrient rich (LB) and nutrient poor (M9) media. Paired, two-tailed T tests were used to test for significance between strains, and growth conditions. Only significant differences are indicated: *, P<0.05. **(E)** Visualisation of PNAG production on KP WT following adherent growth in nutrient rich LB (Top) and nutrient poor M9 minimal media (Bottom). A representative image is shown from 3 (A and B) or 2 (E) biological replicates.

As PNAG is implicated in *Staphylococcal* biofilm formation, we next assessed PNAG production in KP WT following static growth on glass slides, using **Δ***pgaC* as a negative control. Following adherent growth, KP WT produced large extracellular PNAG networks connecting adherent bacteria, while **Δ***pgaC* did not produce detectable PNAG (Figure 1B). We next used enzyme linked immunosorbent assay (ELISA) to obtain a quantitative readout of PNAG production from KP WT and **Δ***pgaC* strains. Probing with the highest concentration of mAb F598 (250 ng/mL) resulted in absorbance readings of 1.2 (450 nm) in KP WT, while binding was negligible in **Δ***pgaC* (Figure 1C). We observed dose-dependent binding of mAb F598 in KP WT confirming that binding was PNAG-specific (Figure 1C). Therefore, we concluded that KP WT produces detectable PNAG *in vitro* but only during adherent growth conditions.

Given that PNAG production in KP WT was dependent on growth conditions, we next assessed the contribution of PNAG to KP biofilm formation. Loss of PNAG did not affect KP biofilm formation in either nutrient poor (M9 minimal media) or nutrient rich (Luria Bertani (LB)) culture media (Figure 1D). Interestingly, PNAG production was markedly reduced in M9, when compared to growth in LB, despite significantly higher biofilm formation (Figure 1D and E). This suggests that PNAG does not contribute to biofilm formation in KP, irrespective of nutrient availability.

### PNAG production is widespread across KP clinical isolates

While the *pga* locus is present in up to 85 % of KP genomes, the detection of PNAG across KP strains has not been reported [8]. Therefore, we investigated if the extracellular PNAG networks observed in KP WT are also seen in a collection of diverse KP clinical isolates. We utilised a panel of 100 KP clinical isolates, collected over a 19-year period (2001-2020) that represents the contemporary diversity of the species [23]. We grew the 100 KP clinical isolates adherently to glass slides, using KP WT and **Δ***pgaC* as controls, and probed them with mAb F598. We detected PNAG production in 91/100 clinical isolates, including isolates belonging to the high risk MLSTs ST11, ST512, ST15, ST101, ST147 and ST307 [24] (Figure 2A and S1A-B). Despite all encoding an intact *pga* locus, we did not detect PNAG on 9 isolates in the collection belonging to ST5445, ST111, ST86, ST45, ST5449, ST380, ST11, ST147 and ST258. As ST258 is a highly disseminated clonal group, we assessed PNAG production in two additional ST258 isolates: KP35 and MKP103. In both strains PNAG was not detected, suggesting that this phenotype is conserved across ST258 isolates (Figure S2).

**Figure 2.**
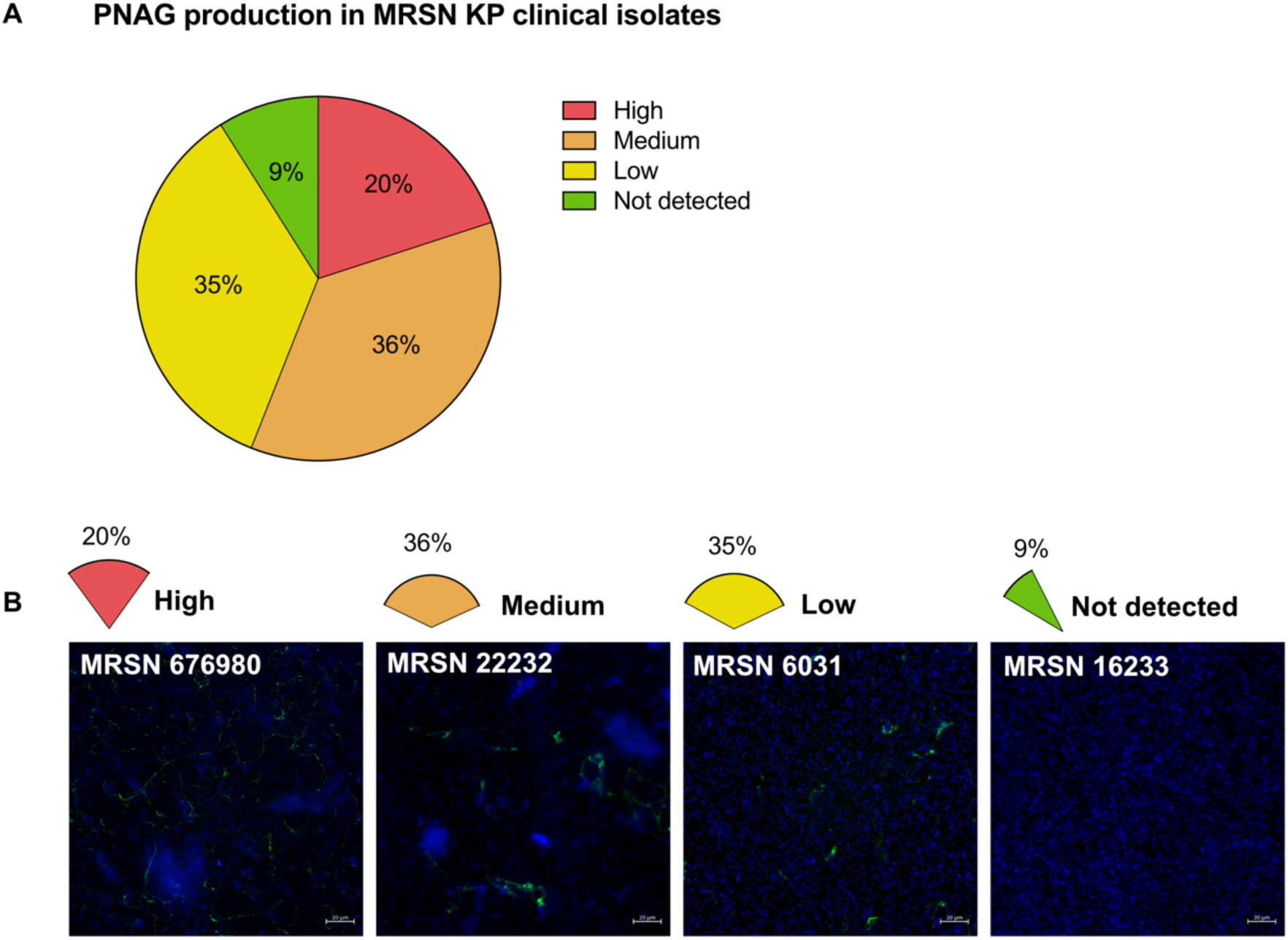
PNAG production is prevalent across KP clinical isolates. **(A)** Prevalence of PNAG production detected on KP clinical isolates of the MRSN during adherent growth on glass slides. PNAG producing phenotypes were assigned by comparing PNAG production on MRSN isolates to KP WT and **Δ***pgaC* where: high > KP WT, medium = KP WT, low < KP WT and not detectable = **Δ***pgaC*. **(B)** Representative images of high, medium, low and not detectable PNAG production across KP clinical isolates. In each case, KP is stained with FM4-64 (blue) and PNAG with mAb F598 (green), and unique MRSN isolate IDs are provided. A representative image across two biological replicates is shown.

By comparing the fluorescence signal from mAb F598 binding, we defined four PNAG-producing phenotypes across the collection: high (>KP WT), medium (=KP WT), low (<KP WT) and no PNAG detectable (=**Δ***pgaC*). Of the 91 strains that produced detectable PNAG, 20, 36 and 35 isolates were characterised as high, medium and low PNAG producers, respectively (Figure 2A-B and S1A-B). All detectable PNAG in the KP clinical isolates formed extracellular networks (Figure 2B). This is consistent with KP PNAG being secreted into the external environment during adherent growth. Taken together, this confirms that the production of extracellular PNAG networks is conserved across KP clinical isolates and validates KP WT as a representative model strain to study PNAG production in KP.

### PNAG does not contribute to KP virulence in a model of severe pulmonary infection

Loss of PNAG production attenuates KP virulence in experimental model of lethal peritonitis in mice; however, the contribution of PNAG to KP virulence during lung infection has not been explored [19]. To address this, we infected BALB/c mice with 250 CFU KP WT and **Δ***pgaC* via intratracheal (IT) intubation, which closely mimics the route of KP infection during ventilator associated pneumonia [25–27]. At 48 h post infection (hpi), we assessed physical readouts of disease pathology, and harvested mouse lungs and blood to quantify bacterial burden and assess the production of pro-inflammatory cytokines (Figure 3A). Mice infected with both KP WT and **Δ***pgaC* had no significant differences in weight loss or wet lung mass index, a good indicator of lung injury encompassing cellular infiltrate and inflammatory oedema (Figure 3B and C). Furthermore, there was no difference in bacterial burdens in the lungs and blood of infected mice (Figure 3D and E). Quantification of the inflammatory cytokines IFNy, CXCL1, TNF, IL6 and GCSF in serum at 48 hpi, revealed no differences between mice infected with KP WT and **Δ***pgaC* (Figure 3F-J). These data confirm that PNAG does not contribute to KP virulence during severe lung infection.

**Figure 3.**
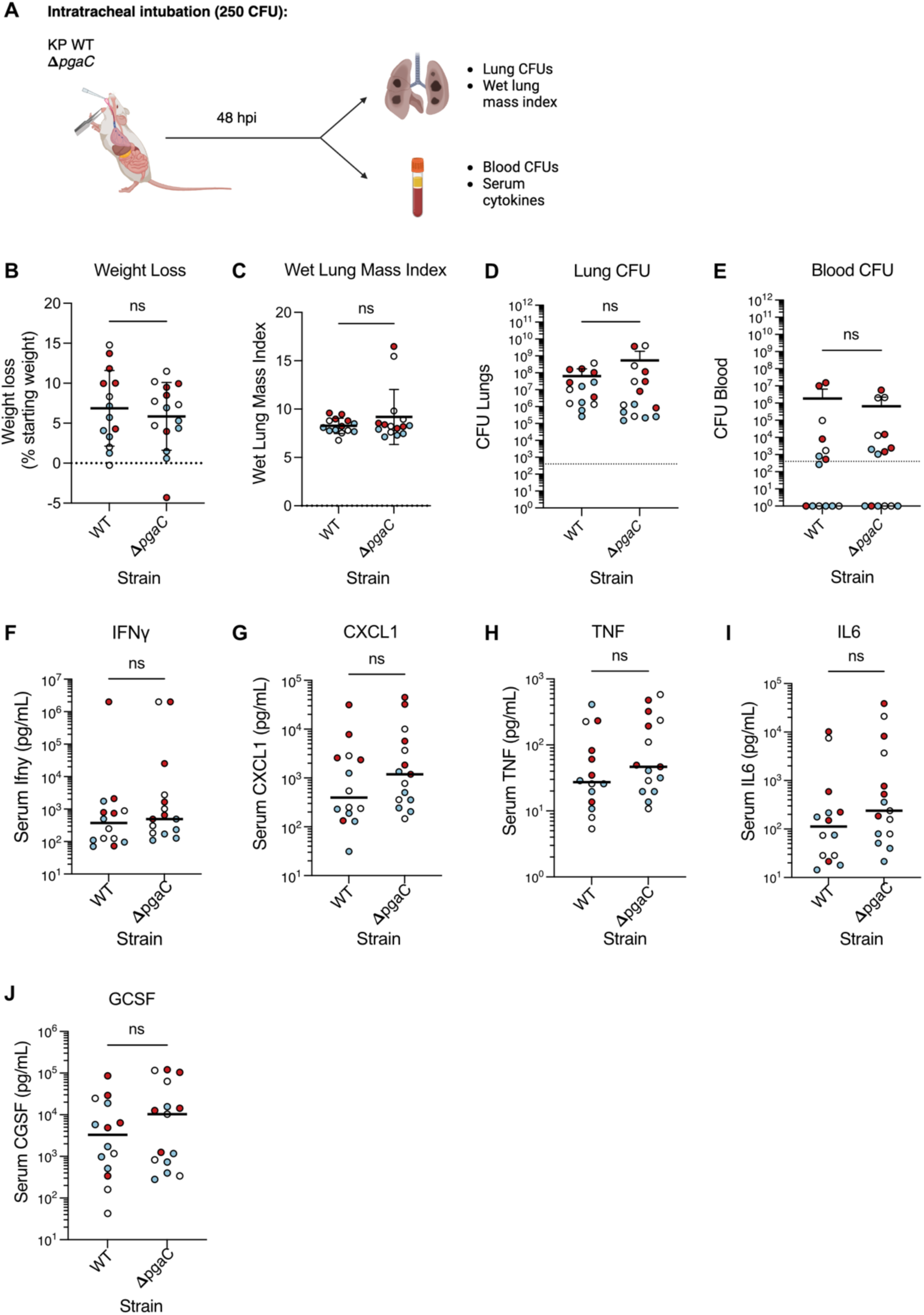
PNAG does not contribute to KP virulence during severe pulmonary infection in mice. **(A)** Simplified model of intratracheal intubation of BALB/c mice with 250 CFU KP WT and **Δ***pgaC* and associated infection readouts 48 hpi. **(B and C)** Physical readouts of weight loss and wet lung mass index (ratio of lung weight to starting body weight) in infected mice 48 hpi. Enumeration of bacterial CFUs in the lungs **(D)** and blood **(E)** of infected mice 48 hpi. **(F-J)** Quantification of inflammatory cytokines detectable in the serum of infected mice at 48 hpi. Graphs show means ±sd (B-E) or median (F-J) across 3 biological replicates; coloured symbols represent data points from the same biological repeat. Statistical significance was assessed using unpaired, two-tailed t tests for normally distributed data (all apart from blood CFUs, serum IFNy and IL6). For non-normally distributed datasets, non-parametric Mann-Whitney tests were used to test for significance; ns, non-significant.

### KP produces cell-associated PNAG during severe pulmonary infection

We next used to immunofluorescence imaging to characterise PNAG production in KP WT in the infected lung. To this end, we utilised a perfusion and inflation protocol to clear the pulmonary vascular tree of red blood cells and maintain lung architecture, which would ordinarily collapse upon removal post-mortem (Figure S3A) [28]. Inflation and simultaneous fixation of perfused lungs with paraformaldehyde (PFA) also acted to preserve PNAG architecture. We then removed the large single left lobe of inflated lungs which was sectioned and stained for histopathological analysis and immunofluorescence imaging. We first analysed H&E stained lung sections to identify areas of pneumonia, characterised by dense alveolar consolidation with immune cell infiltration by sectioning the left lobe of infected lungs in 1 mm increments from the base. In the central region of a representative KP WT-infected lung, we observed dense consolidation and immune cell infiltration into the lung parenchyma at a depth of 4 mm (Figure S3B). There was also a significant burden of disease in the conducting airways (bronchioles) defining bronchopneumonia (Figure 4A and S3B). These disease features were replicated in a representative **Δ***pgaC* infected lung at a depth of 6 mm (Figure S3C). Interestingly, H&E-stained adjacent sections (1mm apart) exhibited minimal features of disease, revealing that KP WT infection was localised to specific areas of the lung, indicative of focal pneumonic process (Figure S3B-C).

**Figure 4.**
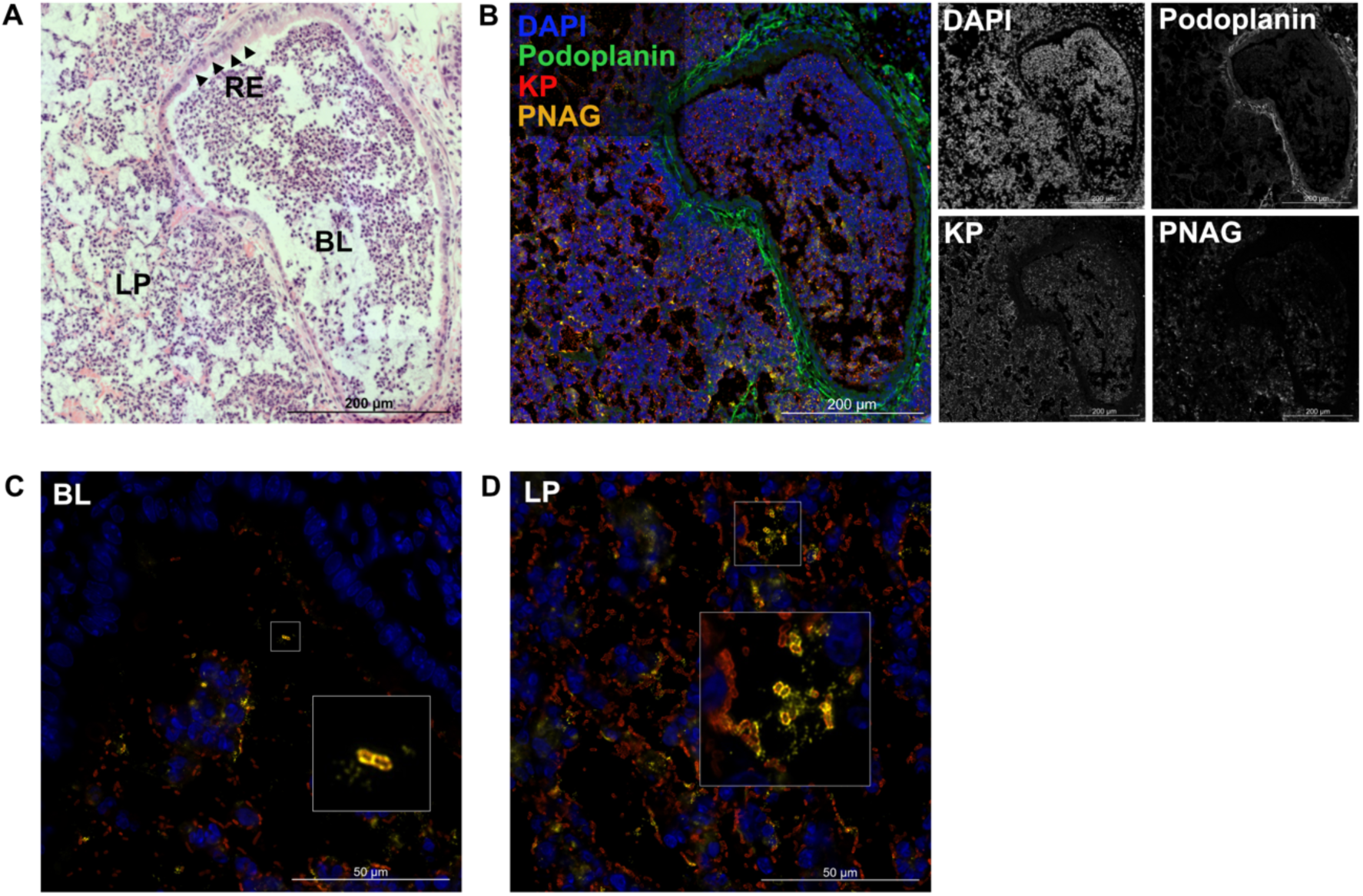
KP undergoes a switch to cell-associated PNAG production during severe pulmonary infection. **(A)** Diseased region of a representative H&E stained KP WT-infected BALB/c lung, with key physiological features labelled, where RE: respiratory epithelium, BL: bronchiole lumen and LP: lung parenchyma. **(B)** The same region of infected lung stained with an antibody mix containing: DAPI, podoplanin, polyclonal KP antibody and mAb F598. Individual channels are shown in black and white, with channel titles representing corresponding colours in the merged image. Magnified images of ICC8001 in the bronchiole lumen **(C)** and lung parenchyma **(D)** demonstrating cell-associated PNAG production in KP WT.(C and D) Only channels for DAPI, polyclonal KP antiserum and mAb F598 are shown.

We next probed areas of disease for KP, PNAG and a structural marker (podoplanin) of type I alveolar epithelial cells (AECs). Staining of the KP WT infected section revealed strong DAPI staining and a concomitant reduction in podoplanin staining in the diseased central portion of the lung, reflective of immune cell recruitment and lung epithelia disruption, respectively (Figure S4A). We also observed strong KP staining in both the alveolar spaces and bronchiole lumen. PNAG staining was observed across the breadth of this KP rich region, spanning both lung parenchyma and bronchioles (Figure 4B and S4A). In comparable regions of **Δ***pgaC*-infected lung we observed the same features with absent PNAG staining (Figure S4B). Taken together, this confirms that KP WT produces PNAG during *in vivo* lung infection in mice.

Upon closer inspection at higher magnification, we observed a new PNAG phenotype in the KP WT infected lung. Together with extracellular networks (often coating alveolar wall structures), as observed in adherent growth in vitro, bacterial cell associated PNAG was present. KP cell-associated PNAG was identified in both the bronchial lumen and regions of dense alveolar consolidation and presents as circumferential staining of individual bacteria (Figure 4C-D). Given we did not observe cell-associated PNAG production during adherent growth *in vitro*, we conclude that this is a feature which may only become apparent in the *in vivo* environment.

### Loss of capsule results in cell-associated PNAG production

The adaptation of KP to the *in vivo* environment is not well understood including the regulation and expression of key virulence factors such as LPS, CPS, type 3 fimbriae and siderophores [12]. Given the switch to a cell-associated PNAG phenotype *in vivo* we hypothesised that there may be a regulatory change in a KP cell wall component underpinning this change. To test this, we engineered a panel of isogenic deletion mutants of prominent KP cell surface molecules, including those involved in capsule synthesis (**Δ***wcaJ*) [29] and regulatory (**Δ***rmpADC*) genes, the LPS O-antigen (**Δ***rfb*), as well as the prominent outer membrane proteins (OMPs), OmpA (**Δ***ompA*) [29] and OmpK36 (**Δ***ompk36*) [27] (Figure 5A). While deletion of *wcaJ* abolishes capsule production, the deletion of *rmpADC* genes results in reduced abundance (Figure S5) [30,31]. We grew the isogenic mutants under adherent growth *in vitro* and examined PNAG localisation by immunofluorescence microscopy. In both mutants with capsular defects (**Δ***wcaJ* and **Δ***rmpADC*), we observed a mixture of extracellular networks and cell-associated PNAG (Figure 5B). No other mutants exhibited this phenotype, and phenocopied KP WT (Figure 5B). This suggests that cell-associated PNAG production in KP during *in vivo* infection may result from reduced expression of the capsule in the lungs.

**Figure 5.**
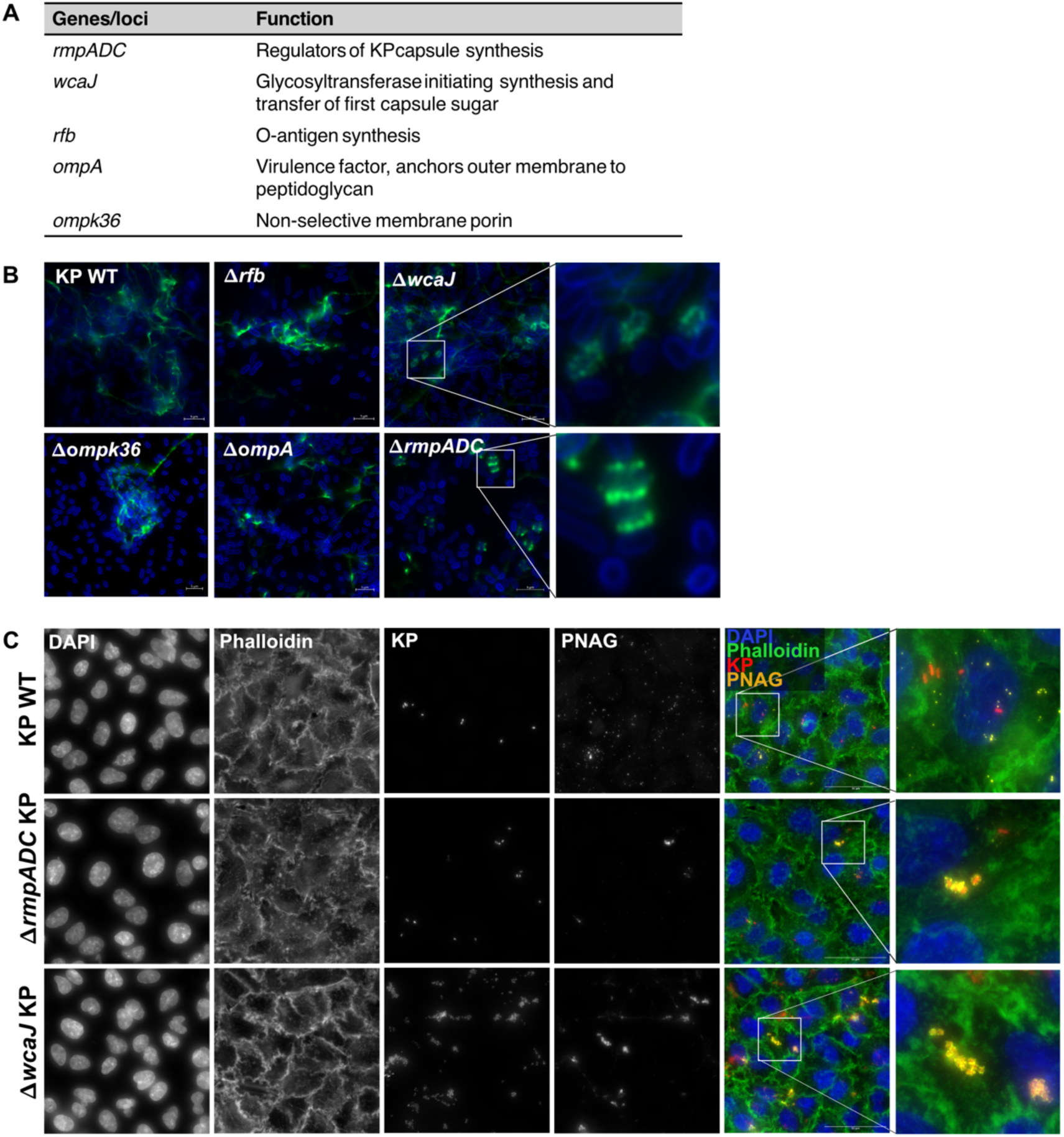
Reduced capsule production on KP WT results in cell-associated PNAG production. **(A)** Panel of isogenic mutants lacking components of the bacterial surface, and each gene/locus function in KP. **(B)** Visualisation of PNAG (mAb F598 in green) production on (FM4-64 in blue) isogenic mutants following adherent growth on glass slides. In each case, a representative merged image across 3 biological replicates is shown. Zoomed images of cell-associated PNAG on **Δ***wcaJ* and **Δ***rmpADC* are shown inset. **(C)** Immunofluorescence staining of PNAG on GFP-expressing KP WT, **Δ***rmpADC* and **Δ***wcaJ* KP during A549 infection (4 hpi, MOI 20). Coloured text above each channel indicates colours in the merged image, with zoomed regions of interest inset in the merged image. In each case, a representative image from 3 biological experiments is shown.

To test this hypothesis, we next compared PNAG localisation on KP WT, **Δ***rmpADC* and **Δ***wcaJ* during *in vitro* infection of A549 cells, a type II AEC line. Infection of A549s with KP WT resulted in poor bacterial attachment, and the production of diffuse extracellular PNAG (Figure 5C). Infection with ***Δ**rmpADC* did not affect bacterial attachment, however, a portion of adherent KP produced cell-associated PNAG (Figure 5C). Meanwhile, loss of capsule on ***Δ**wcaJ* resulted in a marked increase in KP adherence to AECs and a dominant cell-associated PNAG phenotype (Figure 5C). Taken together, this demonstrates that reduced capsule production on KP WT facilitates a switch from extracellular, to cell-associated PNAG production.

## Discussion

PNAG production in KP is incompletely characterised, limiting our ability to evaluate PNAG as a potential protective antigen and therapeutic target for this critical priority organism. Here, we show that KP produces large extracellular networks of PNAG during adherent growth conditions, a phenotype conserved across a panel of KP clinical isolates, spanning multiple high risk MLSTs [24]. However, we were unable to detect PNAG production on 3 isolates belonging to the ST258 clonal group. This differs from ST11 and ST147 isolates which exhibited variation in PNAG production suggesting that this is a conserved phenotype amongst ST258 isolates. Therefore, further investigation is required to determine if this phenotype is associated with the success of the ST258 clonal group as a human pathogen. Unlike other PNAG-producing bacterial pathogens, we were unable to detect PNAG on KP during growth in liquid culture, suggesting that KP PNAG production is dependent on growth conditions and tightly regulated. Indeed, all the KP clinical isolates that did not produce detectable PNAG during adherent growth (9%) contain an intact *pga* locus. While PNAG was highly produced during adherent growth conditions, we confirm that PNAG does not contribute to KP biofilm formation and is markedly reduced during strong biofilm-inducing conditions during growth in minimal media.

We also assessed PNAG production on KP during an *in vivo* model of severe pulmonary infection, a context in which understanding expression is clinically relevant as an antigenic target. While PNAG production did not contribute to virulence in this context, we detected robust PNAG production on KP WT in the lungs of infected mice. Crucially, we found that the *in vivo* lung environment induced a novel PNAG phenotype in KP, switching from exclusively extracellular networks during *in vitro* assays, to a combination of extracellular networks and cell-associated PNAG production in the infected lung. This phenotype was observed in both the alveolar air spaces and bronchiole lumen. This highlights that PNAG production is a dynamic process in KP, changing markedly between *in vitro* and *in vivo* environments. This also suggests that anti-PNAG based therapies could exert dual functionality against KP infections, supporting PNAG as an attractive KP antigen.

Traditionally, attractive bacterial antigenic targets also contribute to virulence, as their requirement for infection reduces the likelihood of antigenic escape. Furthermore, protective antibodies can function to both opsonise and kill invading pathogens directly and block the natural course of infection [32]. However, we demonstrate that PNAG is dispensable for KP lung infection. This differs from *in vivo* models of KP peritonitis, during which loss of PNAG reduces KP lethality in mice [19]. Furthermore, loss of PNAG from hvKP also reduces KP colonisation of the GI tract and dissemination into other organs following oral inoculation [19]. This suggests that the contribution of PNAG to KP virulence is context dependent, reducing the overall likelihood of antigenic escape.

We also present evidence that the switch from extracellular to cell-associated PNAG production during *in vivo* lung infection occurs via a novel interaction between PNAG and capsule on the KP surface. During *in vitro* adherent growth on glass slides, ***Δ**wcaJ* and ***Δ**rmpADC* KP produced predominantly extracellular PNAG networks, while a proportion of cells exhibited a cell-associated PNAG phenotype. This could be explained by capsule and PNAG interacting on the KP surface, or simply, reduced capsule production improving antibody accessibility to PNAG in KP. However, in the context of *in vitro* infection, the predominant PNAG phenotype on KP capsule mutants changed to cell-associated PNAG production, with negligible extracellular network formation. We also found that the prevalence of cell-associated PNAG negatively correlated with capsule production. This is consistent with there being crosstalk between capsule and PNAG on the KP surface, whereby, reduced capsule production facilitates a switch to cell-associated PNAG production in KP.

While the capsular polysaccharide is a characteristic feature of KP strains, there is growing evidence that capsule production across KP clinical isolates can be downregulated during infection. A longitudinal study of KP clinical isolates in China found that a proportion of isolates exhibited an acapsular phenotype, caused by the insertion of selfish DNA elements into the initial glycosyltransferase genes of the capsule synthesis locus [33]. Furthermore, KP bloodstream and urinary tract infection isolates have been found to carry mutations in capsule biosynthesis genes, conferring reduced capsule production [34,35]. Interestingly, in each of these instances, reduced capsule production was associated with improved bacterial fitness, conferring improved resistance to serum bactericidal activity but greater complement mediated killing (32-34). Consistent with this, we demonstrate that loss of capsule in KP results in a marked increase in adherence to AECs. However, there is clearly a fine balance, as deletion of both *wcaJ* and *rmpADC* from hvKP strains results in severe attenuation during *in vivo* models of KP pneumonia via intranasal inoculation [30,31]. With this in mind, we propose that during severe pulmonary infection in mice, KP capsule production is down-regulated, resulting in improved KP adherence to AECs, and a switch to cell-associated PNAG production. Further characterisation of KP capsule production during *in vivo* lung infection would be useful to further characterise the interaction between PNAG and capsule on KP.

Currently, KP capsule and O-antigen represent attractive antigens, and are prominent targets in the design of novel therapeutics against KP infection. However, the serological diversity of both O-antigen and capsule types across KP represents a hurdle to the development of broadly effective treatments [36]. By contrast, PNAG is structurally conserved across KP strains and other bacteria. While interactions between KP capsule and LPS have been described, the relationship between PNAG and other surface polysaccharides on KP have not been reported [37]. Our proposed interaction between PNAG and capsule on KP raises questions as to how bacterial surface antigens interact during infection, and thus, which bacterial polysaccharides represent the most attractive targets. In the case of KP, which seems to exhibit a dynamic surface polysaccharide composition during infection, a multivalent vaccine comprising components of KP capsule, LPS and PNAG, may represent the most effective design option. Overall, our results reinforce PNAG as an attractive bacterial antigen, and prompt further characterisation of PNAG production across priority bacterial pathogens.

## Materials and methods

### Bacterial strains and growth conditions

The strains used in this study are listed in Table S1. All KP strains were grown overnight (O/N) in LB broth at 37℃ under agitation (200 rpm). For ICC8001 strains, media was supplemented with streptomycin (10 µg/mL), unless otherwise stated.

Plasmids and primers used to generate isogenic strains are listed in Tables S2 and S3, respectively. To generate isogenic ORF deletion mutants, the 5’ and 3’ 500 bp flanking genes of interest were assembled into linearised pSEVA612RS, a derivative of pSEVA612S with a flipped I Sce-I site to reduce non-specific recombination, using Gibson assembly. Gene deletions were then introduced into ICC8001 using a 2-step recombination protocol as described previously [38]. Successful deletion was confirmed by PCR and Sanger sequencing.

### Immunofluorescence staining of PNAG *in vitro*

The antibodies used in this study are listed in Table S4. To detect KP PNAG during growth in liquid culture, a modified staining in suspension protocol was used. Briefly, KP O/N cultures were diluted 1:100 (v/v) into PBS and centrifuged at 5000 x g for 5 minutes. Cell pellets were fixed in 1 mL 4% PFA in PBS for 20 minutes at room temperature (RT). Cells were washed twice in 1 mL sterile PBS and resuspended in 100 µL mAb F598 (a kind gift from GSK, Siena, Italy) (2 µg/mL) in 2% bovine serum albumin (BSA) diluted in PBS and incubated on a spinning wheel for 2.5 hours at RT. Cells were centrifuged at 10000 x g for 3 minutes and washed 3 times in PBS. Cell pellets were resuspended in 100 µL fluorescently labelled goat anti-human IgG antibody (AlexaFluor 488, #A-11013, ThermoFisher Scientific) antibody diluted in 2% BSA-PBS and incubated on a rocker for 1 hour at RT in the dark. Cells were washed 3 more times in sterile PBS and resuspended in 100 µL FM4-64 (#F34653, ThermoFisher Scientific) diluted in de-ionised water and incubated for 20 minutes at RT in the dark. Stained cells were washed and resuspended in 1 mL sterile PBS. To image, 2.5 µL stained cells were spotted onto a glass slide, covered with an agarose pad (1% agarose in de-ionised water) and secured in place with a coverslip. Cells were imaged immediately using Zeiss Axio Observer Z1 microscope.

To detect PNAG during adherent growth, KP O/N cultures were diluted 1:1000 (v/v) into fresh LB, and 400 µL added to wells on 8-well IBIDI glass bottomed slides (IBIDI). Slides were incubated at 37℃ statically for 2 h, after which media was aspirated and replaced with 400 µL fresh LB and slides incubated at 37℃ statically for 24 h. Wells were washed in sterile PBS and fixed for 20 mins in 4% PFA-PBS. To detect PNAG, wells were probed with mAb F598 (2 µg/mL) in 2% BSA-PBS for 1 h at RT. Wells were washed in PBS 3 times then probed with fluorescently labelled goat anti-human IgG antibodies (1:200) diluted in in 2% BSA-PBS for 1 h at RT in the dark. Wells were washed 3 times in PBS, then incubated with FM4-64 for 20 minutes at RT in the dark. Wells were washed a final time, resuspended in 400 uL PBS and imaged immediately.

### KP whole cell ELISA

KP O/N cultures were diluted 1:1000 (v/v) into fresh LB and added to wells in a 96-well tissue culture treated plate (Avantor). Plates were incubated at 37℃ statically for 2 h, when media was replaced with 200 uL fresh LB, and plates returned to 37℃ for 24 h. After 24 h, media was aspirated, wells were washed in PBS + 0.05% Tween-20 (PBST) and fixed in 4% PFA-PBS for 20 minutes. Wells were washed 3 times in PBST and incubated with mAb F598 (1 µg/mL for top standard and then serially diluted 2-fold down the plate) in 1% BSA-PBST O/Nt at 4℃. Wells were triple washed in PBST and incubated with goat anti-human IgG conjugated to streptavidin horse radish peroxidase (HRP) (#A18811, ThermoFisher Scientic) for 1 h at RT under agitation. Wells were washed 3 times in PBST and incubated with 3,3’,5,5’-Tetramethylbenzidine (TMB, #N301, ThermoFisher Scientific) in the dark. Stop solution was added to wells when a clear gradient was visible across mAb F598 dilutions. Absorbance readings at 450 and 540 nm were taken immediately using a FLUOstar Omega plate reader.

### Biofilm assay

KP O/N cultures were diluted 1:1000 (v/v) into fresh LB or M9 minimal media + 0.4 % glucose and added (200 µL) to wells in a 96-well tissue-culture treated plate (Avantor). Plates were incubated statically at 37℃ O/N. Wells were washed 3 times H_2_O then incubated with 300 uL crystal violet (0.1% w/v) for 20 mins at RT on a rocker. Wells were washed 3 more times in H_2_O and left to dry O/N at RT. Crystal violet-stained wells were resuspended in 200 uL de-stain solution (30% acetone w/v) and transferred to a fresh 96-well plate. Absorbance readings were taken at 595 nm using a FLUOstar Omega plate reader.

### Animal experiments

Pathogen-free female BALB/c 18- to 20-g female mice were purchased from Charles River Laboratories and housed in groups of 5 in individually HEPA-filtered cages with bedding, nesting, and free access to food and water. The temperature, humidity and light cycles were kept within the UK Home Office code of practice, with the temperature between 20 and 24 °C, the room humidity at 45–65% and a 12 h/12 h light cycle with a 30-min dawn and dusk period to provide a gradual change. Mouse experiments were performed in accordance with the Animals Scientific Procedures Act of 1986 [39] and UK Home Office guidelines and were approved by the Imperial College Animal Welfare and Ethical Review Body. Experiments were designed in agreement with the ARRIVE guidelines [40] for the reporting and execution of animal experiments, including sample randomization and blinding.

In each case, KP inoculum was prepared by diluting KP O/N cultures in PBS to a final concentration of 250 CFU/50 µL. For infections, mice were placed under anaesthesia via IP administration of a mixture of ketamine (80 mg/kg) and medetomidine (0.8 mg\kg) and inoculated via intratracheal intubation, as previously described (24-26).

At 48 hpi, terminal anaesthesia was induced in mice via IP injection of ketamine (100 mg/kg) and medetomidine (1 mg\kg). Once mice had reached deep anaesthesia, they were placed in dorsal recumbency, and blood was collected directly from the heart via cardiac puncture. A portion of blood was mixed immediately with an anti-coagulant (1:10), which was subsequently serially diluted and plated onto rifampicin (50 mcg/mL) LB agar plates to enumerate CFUs. Remaining blood was collected into microtainer serum tubes and centrifuged at 16000 x g to collect serum for subsequent cytokine analysis. For CFU counts, lungs were removed from the thorax postmortem and collected in 3 mL DPBS and homogenised. Lung homogenate was then serially diluted and plated onto rifampicin (50 mcg/mL) LB agar plates to enumerate lung CFUs.

To quantify serum cytokines, a multiplexed beads-based assay was used (LEGENDplex). Briefly, serum samples and detection beads were mixed in a 96-well V-bottomed plate, and incubated for 2 h at RT, shaking (800 rpm). The plate was centrifuged at 500 x g for 5 minutes, then wells were washed and incubated with detection antibody for 1 h at RT, shaking (800 rpm). Streptavidin-phycoerythrin (SA-PE) was added to wells, and plates incubated for a further 30 minutes at RT, shaking (800 rpm). Wells were washed, resuspended in 200 µL wash buffer and transferred to FACS tubes pre-filled with 100 µL wash buffer to reach a final volume of 300 µL. Samples were read immediately using FACScalibur flow cytometer to detect a custom panel of cytokines. Analysis was carried out using LEGENplex data analysis software.

For *in vivo* detection of PNAG on KP, mice were infected via IT intubation as described above, and euthanised 48 hpi via IP injection of pentobarbitone (200 uL). Death was confirmed by cutting the femoral artery. The mouse was placed in dorsal recumbency, and the anterior fur and skin removed. The abdomen was then opened via midline incision, and the bowel removed. The descending aorta and inferior vena cava were cut and the right ventricle was perfused with 10 mL PBS. The mouse was then intubated, and sutures were tied around the end of the intubation canula. The lungs were inflated with 4 % PFA-PBS to 20 cm H_2_O, at which point the intubation catheter was removed, and the sutures fastened to prevent leakage of the fixative. The entire pulmonary tree was removed and immediately moved to 4 % PFA-PBS 2 h at RT then submerged in 70 % ethanol for 24 h at 4 ℃.

### Immunofluorescence staining lung and liver sections

Following fixation, the left lung lobe was paraffin embedded and sectioned. Sections were taken at 1mm increments and hematoxylin and eosin stained (H & E) stained until a depth at which severe disease, defined as alveolar wall thickening and immune cell infiltration, was visible. Once this depth was reached, adjacent unstained sections were taken to be used for immunofluorescence staining.

Unstained lung sections were dewaxed and re-hydrated in ethanol, as described previously [41]. Antigen retrieval was performed by heating sections in de-masking solution (0.3% trisodium citrate-0.05% Tween 20 in distilled H_2_O). Heated sections were left to cool at RT, circled in wax pen, and blocked in PBS + 0.05 % Tween-20 + 0.1 % saponin (PBSTS) for 1 h at RT. Lung sections were probed with the following primary antibody mix: polyclonal KP antibody (1:100, #ab20947, Abcam), mAb F598 (2 ug/mL) and anti-podoplanin antibody (1:100, # ab256559, Abcam) diluted in PBSTS and incubated at 4℃ O/N. The next day, sections were washed twice in PBSTS for 10 minutes on a rocker at RT. Sections were then probed with the following secondary antibody mix: DAPI (1:1000, #D3571, ThermoFisher Scientific), donkey anti-rabbit IgG rhodamine red-X (RRX) (1:100, # 711-295-152, Jackson Immunoresearch), donkey anti-human IgG AF647 (1:100, #709-605-149, Jackson Immunoresearch), donkey anti-rat IgG AF488 (1:100, #712-546-150, Jackson Immunoresearch) diluted in PBSTS for 1 h at RT in the dark. Sections were washed twice in PBSTS and left to dry at RT in the dark. Cover slips were mounted using Prolong Gold antifade mounting media (#P36930, ThermoFisher Scientitifc), and left to harden O/N at RT in the dark. Stained sections were imaged the next day using a Zeiss Axio Observer Z1 microscope.

### *In vitro* infections

A549 cells were maintained Dulbecco Modified Eagle’s Medium (DMEM, #D6546 Sigma) supplemented with 1x GlutaMax (#35050061, ThermoFisher Scientific) and 10% Fetal Bovine Serum (#F9665, Sigma) without antibiotics under static conditions at 37℃, 5% CO_2_.

Cells were seeded (2×10^5^ cells/well) in 400 µL fully supplemented DMEM into tissue culture treated, glass bottomed IBIDI slides and incubated O/N at 37℃, 5% CO_2_. On the day of infection, media was replaced with unsupplemented, low glucose DMEM (#D5546, Sigma). Inoculum was prepared by diluting KP O/N cultures in unsupplemented DMEM to obtain a multiplicity of infection (MOI) of 20 in 20 uL, which was added to wells. Infected slides were centrifuged at 700 x g for 10 mins to synchronise infection and KP inoculum was serially diluted and spotted onto LB agar plates to validate MOIs.

For immunofluorescence microscopy, media was aspirated from infected wells after 4 hours to stop the infection. Wells were washed 3 times in PBS and fixed in 4% PFA-PBS for 20 minutes at RT. Wells were washed 3 times and resuspended in 1 mL PBS. If staining was to be carried out on a different day, wells were stored in the dark at 4℃ at this stage. For staining, wells were quenched with 50 mM ammonium chloride (NH_4_Cl) in PBS for 10 mins at RT, washed once in PBS and permeabilised in 0.1% Triton X-100 in PBS for 10 mins at RT. Once permeabilised, wells were washed 3 times and blocked in 3% BSA-PBS for 20 minutes. Wells were probed with mAb F598 (2 µg/mL) in 2% BSA-PBS for 1 h at RT. Wells were washed 3 more times in PBS and incubated with an antibody mix containing: DAPI (1:1000) (cell nuclei); Phalloidin iFluor 647 (1:100, #23127-AAT, Stratech) (F-actin) and goat anti-human IgG Alexa555 (1:200, #A-21433, ThermoFisher Scientific) for 1 h at RT in the dark. Wells were washed 3 times in sterile PBS, resuspended in 400 µL sterile PBS and imaged immediately using Zeiss Axio Observer Z1 microscope. Images were taken as Z stacks and merged using maximum intensity projection (ImageJ).

### Statistical analysis

All data was analysed in Graphpad Prism 10. All ELISA conditions were performed in triplicate and the mean absorbance calculated to represent each biological replicate. For comparisons between ICC8001 and **Δ***pgaC,* paired and unpaired two-tailed t tests were used to test for the significance of normally distributed data, and Mann-Whitney tests for data that was not normally distributed.

## Acknowledgements

*Klebsiella pneumoniae* MRSN Diversity Panel, NR-55604 was provided by the MultidrugResistant Organism Repository and Surveillance Network (MRSN) at the Walter Reed Army Institute of Research (WARIR), Silver Spring USA through BEI Resources, NIAID, NIH. We would also like to thank Professor Alice Prince for providing KP35 and Tania Wong for providing MKP103.

This work was sponsored by a BBSRC and GlaxoSmithKline (GSK) studentship to JB.

## Supplementary Figures

**Figure S1.**
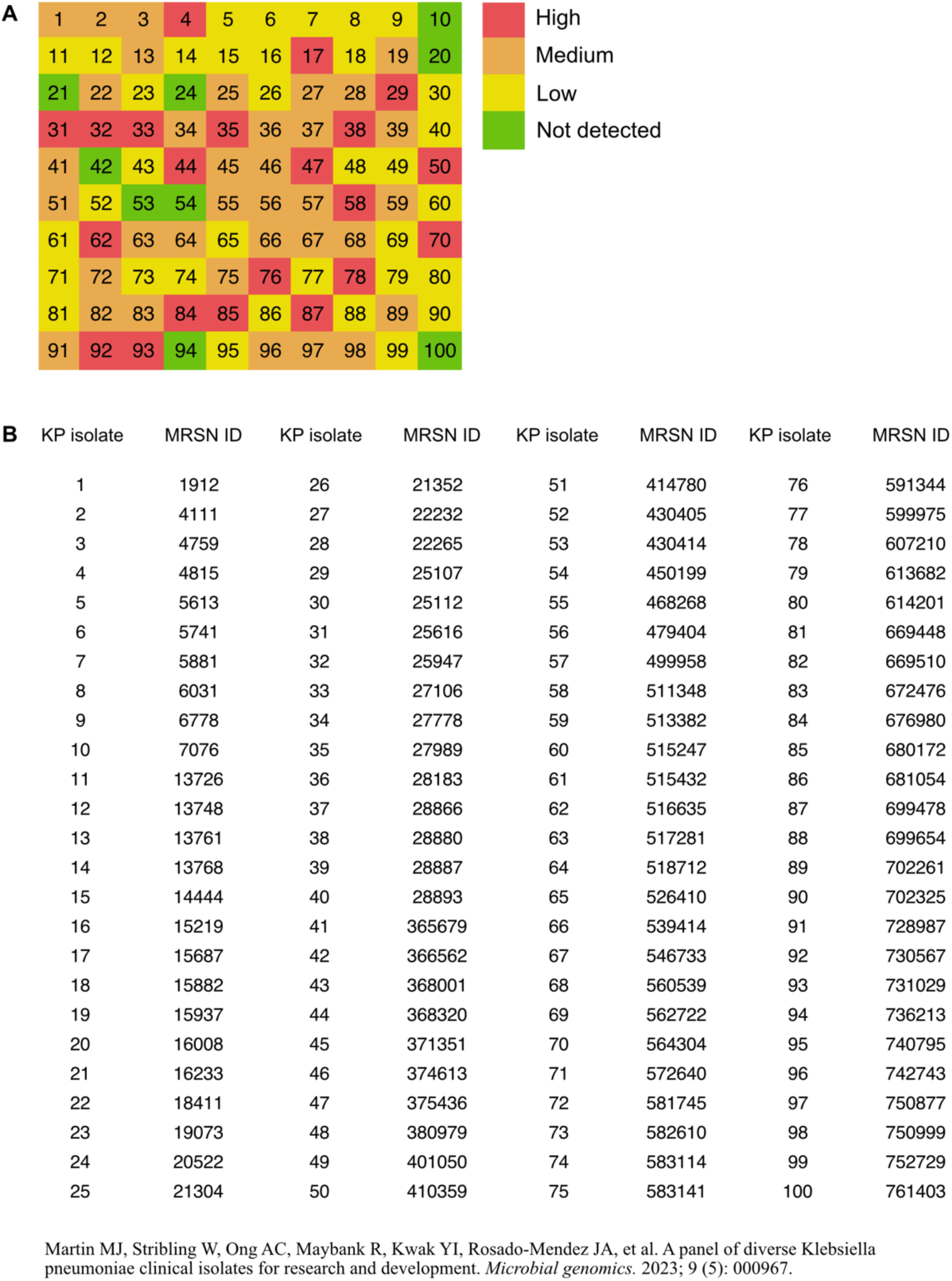
PNAG production across MRSN KP clinical isolates. **(A)** Heat-map showing PNAG producing phenotype of KP isolates in the MRSN collection. The PNAG phenotype was characterised using immunofluorescence microscopy, and comparing to KP WT and **Δ***pgaC,* where high > KP WT, medium = KP WT, low < KP WT and not detected = **Δ***pgaC.* Data is reflective of two biological repeats. **(B)** MRSN identification numbers of the 100 KP clinical isolates, with the associated reference.

**Figure S2.**
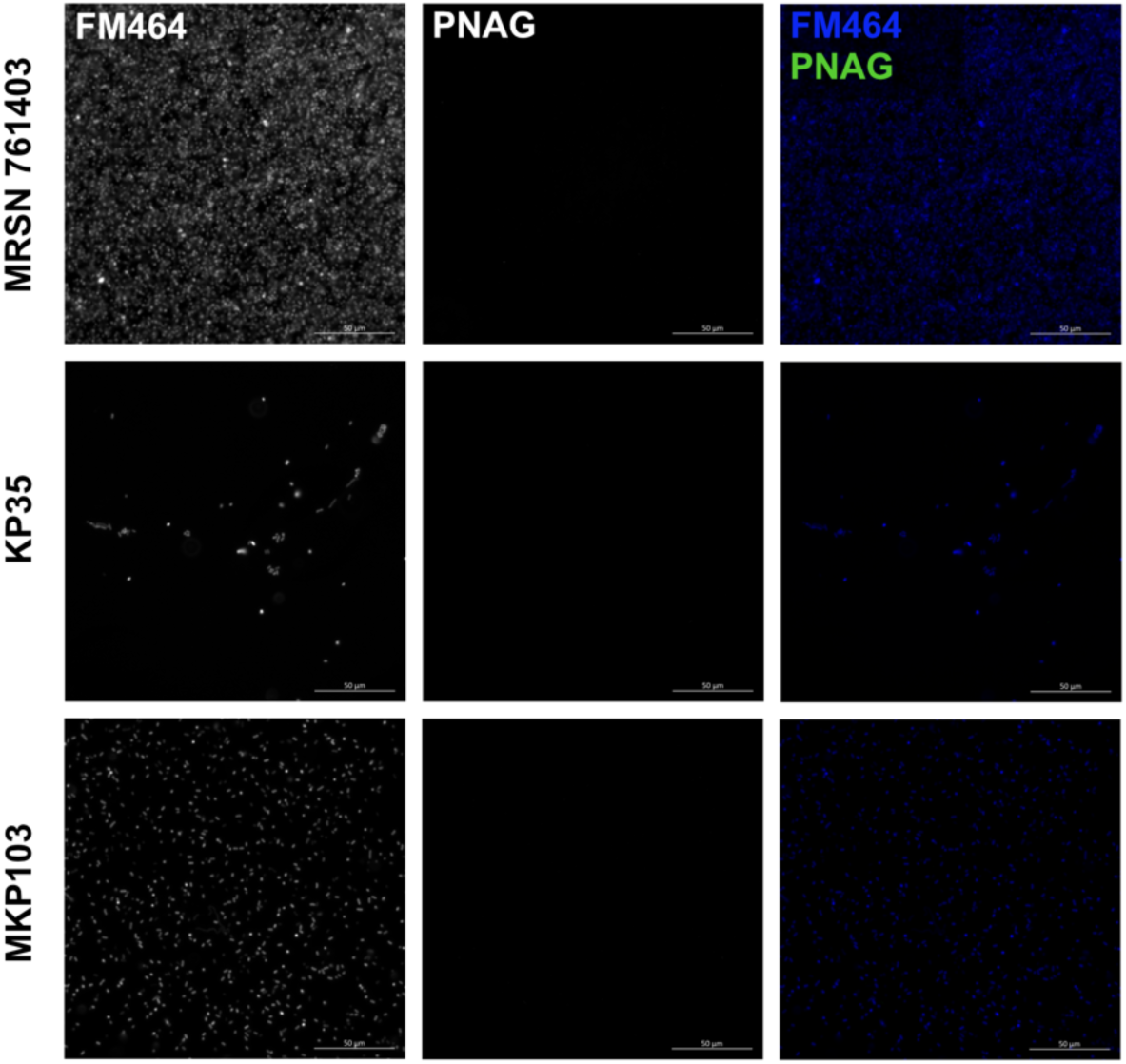
KP ST258 isolates do not produce PNAG during adherent growth conditions. Visualisation of PNAG on ST258 isolates (MRSN761403, KP35 and MKP103) during adherent growth on glass slides. A representative image from 2 biological replicates is shown.

**Figure S3.**
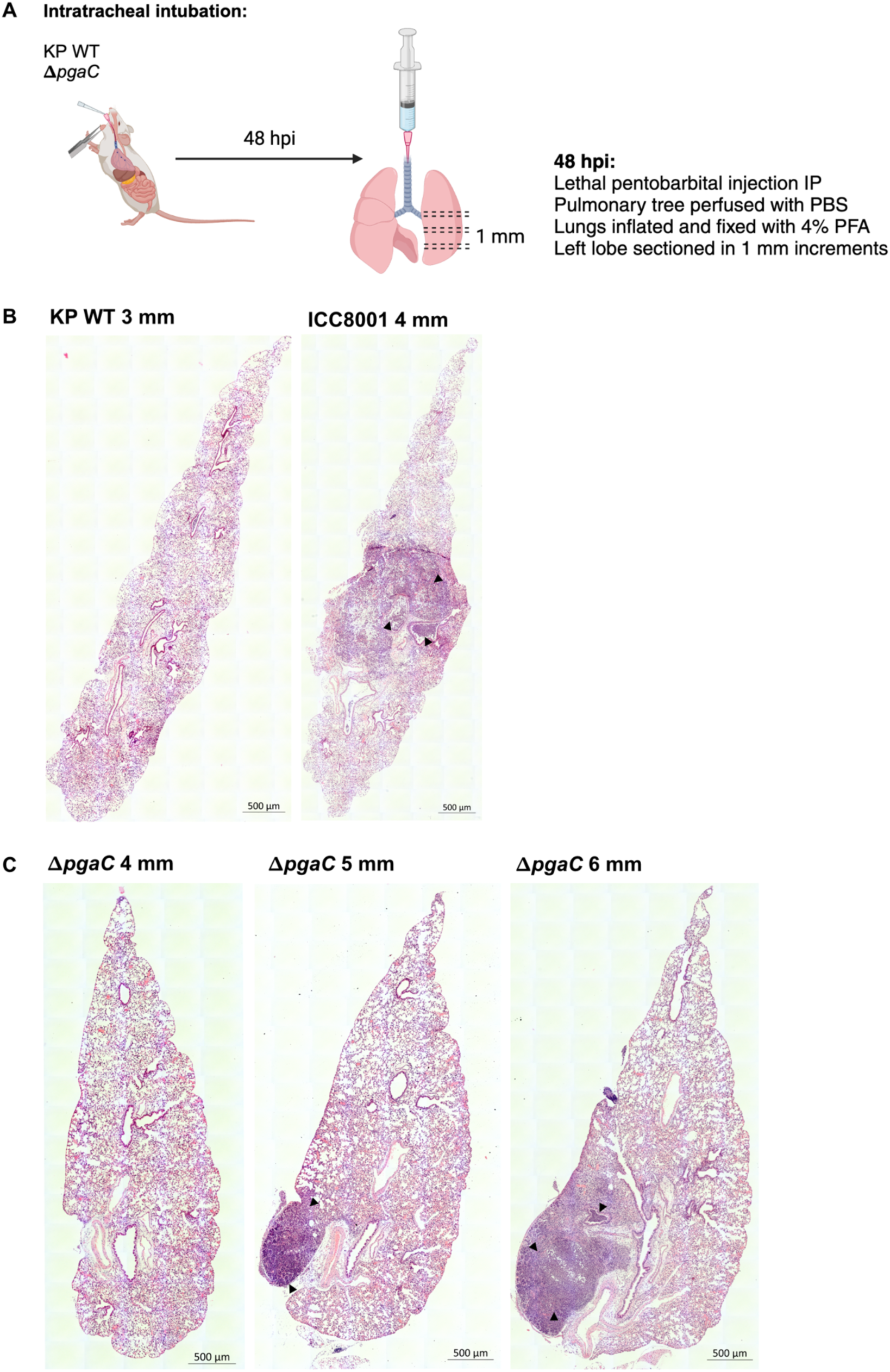
KP WT causes focal pneumonia in BALB/c mice. **(A)** Simplified overview of inflation and perfusion protocol used to preserve lung architecture prior to sectioning and imaging. H&E stained sections of the left lung lobe of a representative **(B)** KP WT and **(C) Δ***pgaC* infected BALB/c mice at 48 hpi. In each case sections exhibiting severe disease pathology, defined as alveolar consolidation with immune cell infiltration, was observed (4 mm in KP WT and 6 mm in **Δ***pgaC*). Adjacent sections, taken 1 mM apart, are also shown to demonstrate the localised nature of KP in the lung. Black arrowheads point to regions of immune cell infiltration into both the lung parenchyma and bronchiole lumen.

**Figure S4.**
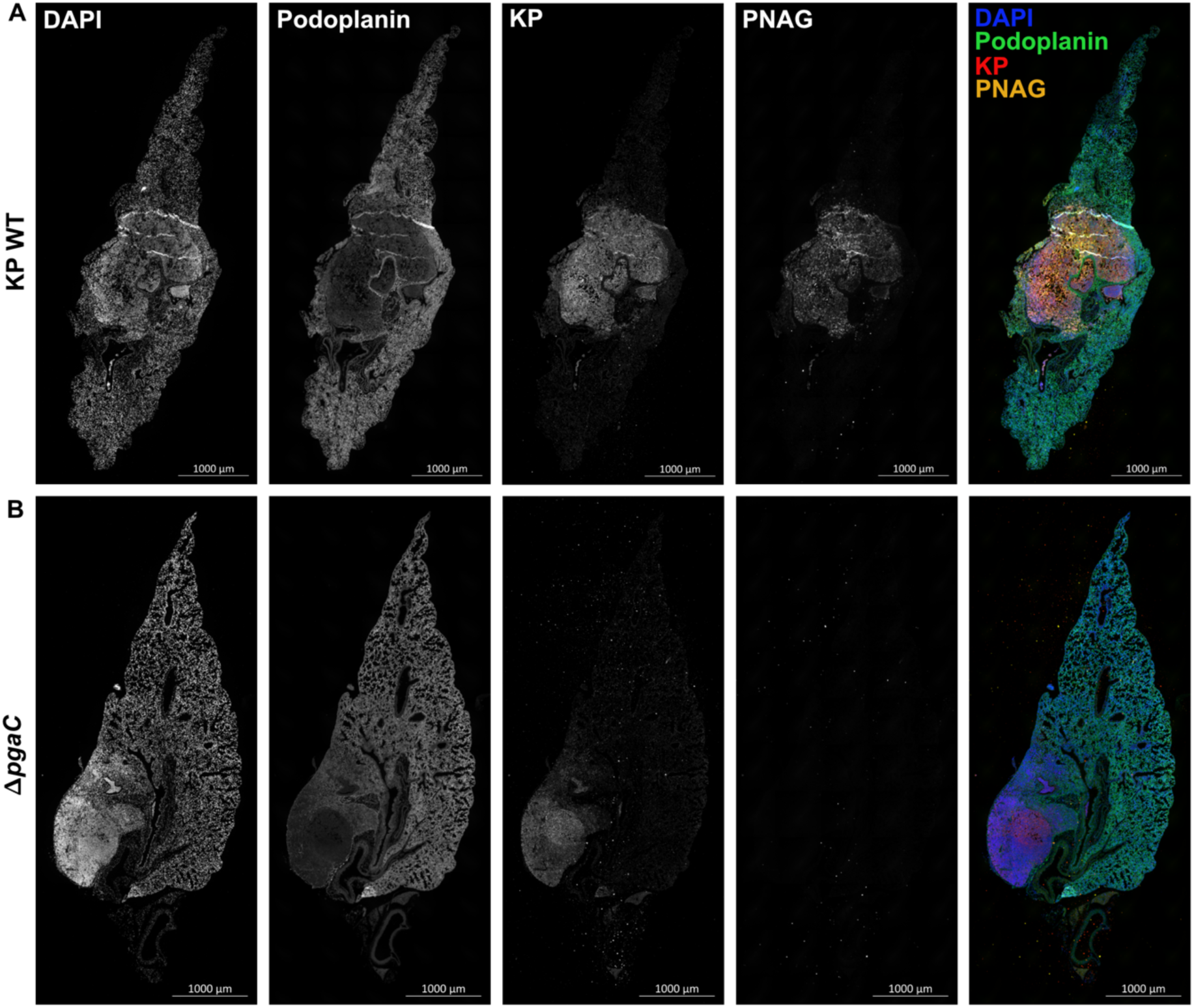
KP produces PNAG during *in vivo* lung infection in BALB/c mice. Immunofluorescence images of KP WT **(A)** and **Δ***pgaC* KP **(B)** infected lung sections exhibiting severe disease pathology 48 hpi (sections identified in Figure S3B and C). In each case, individual channels are shown in black and white alongside a coloured merge, with colours matching channel headings.

**Figure S5.**
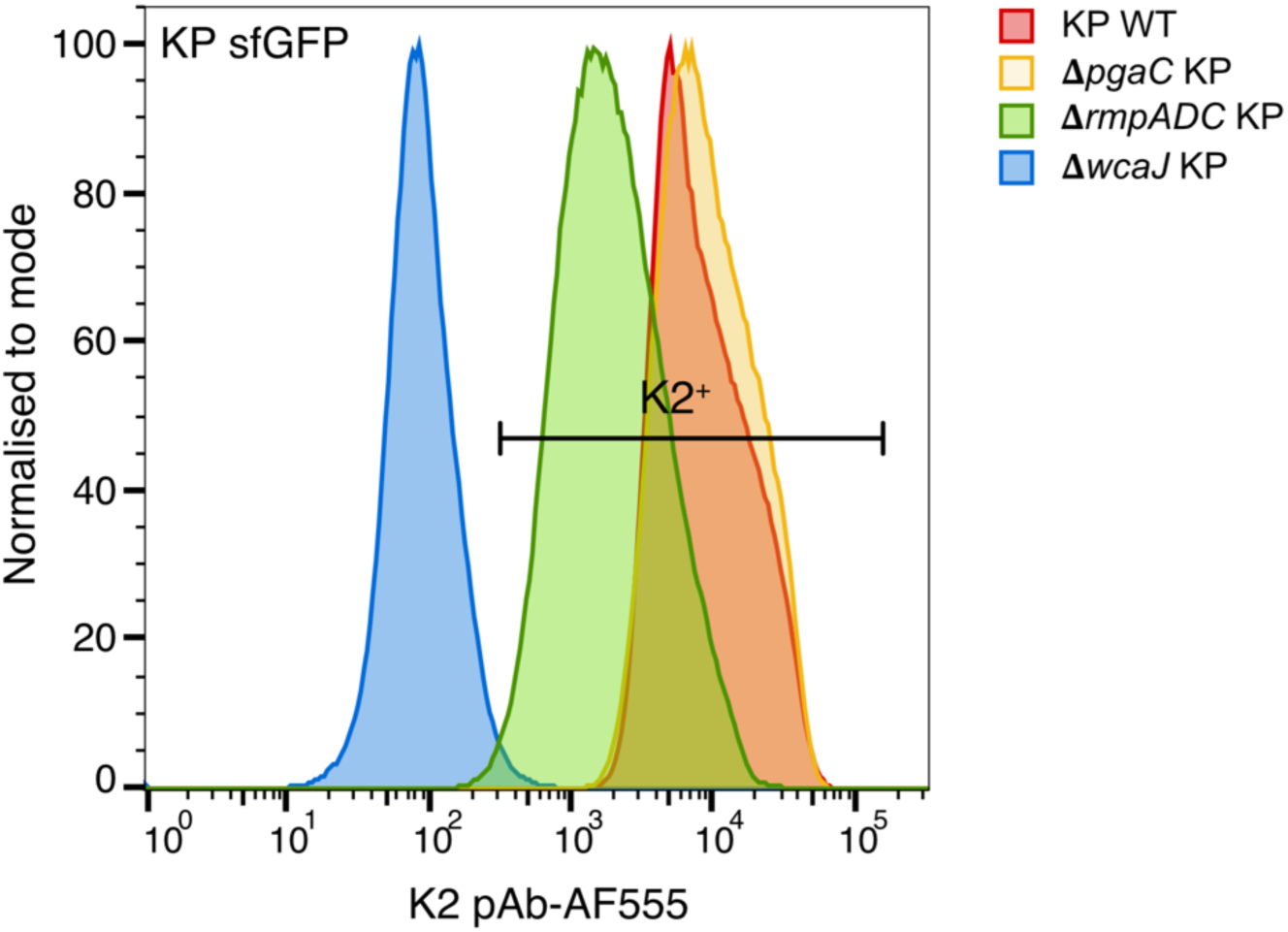
Quantification of capsule production on Δ*wcaJ* and Δ*rmpADC* KP. Flow cytometric analysis of K2 capsule production on fluorescently tagged WT, **Δ***pgaC,* **Δ***rmpADC and* **Δ***wcaJ* KP following overnight growth under agitation. Data is presented as percentages of the maximum intensity.

## Supplementary Tables

**Table S1.** Bacterial strains used in this study.

| Bacterial strains |  | Source |
| --- | --- | --- |
| KP ICC8001 strains | WT | Wong et al 2019 |
|  | WT pUltra-sfGFP | This study |
| | $\Delta pgaC$ | This study |
| | $\Delta pgaC$ pUltra-sfGFP | This study |
| | $\Delta rmpADC$ | This study |
| | $\Delta rmpADC$ pUltra-sfGFP | This study |
| | $\Delta wcaJ$ | Low et al 2022 |
| | $\Delta wcaJ$ pUltra-sfGFP | This study |
| | $\Delta rfb$ | This study |
| | $\Delta ompA$ | Low et al 2022 |
| | $\Delta ompK36$ | Wong et al 2019 |
| KP MRSN isolates |  | Martin et al 2023 |
| *E. coli* strains | *E. coli* CC118λpir | Lab collection |
|  | *E. coli* pRK2013 | Figurski and Helinski 1979 |

**Table S2.** Plasmids used in this study.

| Plasmid | Resistance | Source |
| --- | --- | --- |
| pSEVA612SR | Gentamicin | This study |
| pACBSR | Streptomycin | Ruano-Gallego et al 2015 |
| pUltra-sfGFP | Gentamicin | Gift from Despoina Mavridou |
| pSEVA612SR Δ*pgaC* | Gentamicin | This study |
| pSEVA612SR Δ*rmpA-C* | Gentamicin | This study |
| pSEVA612SR Δ*rfb* | Gentamicin | This study |

**Table S3.**
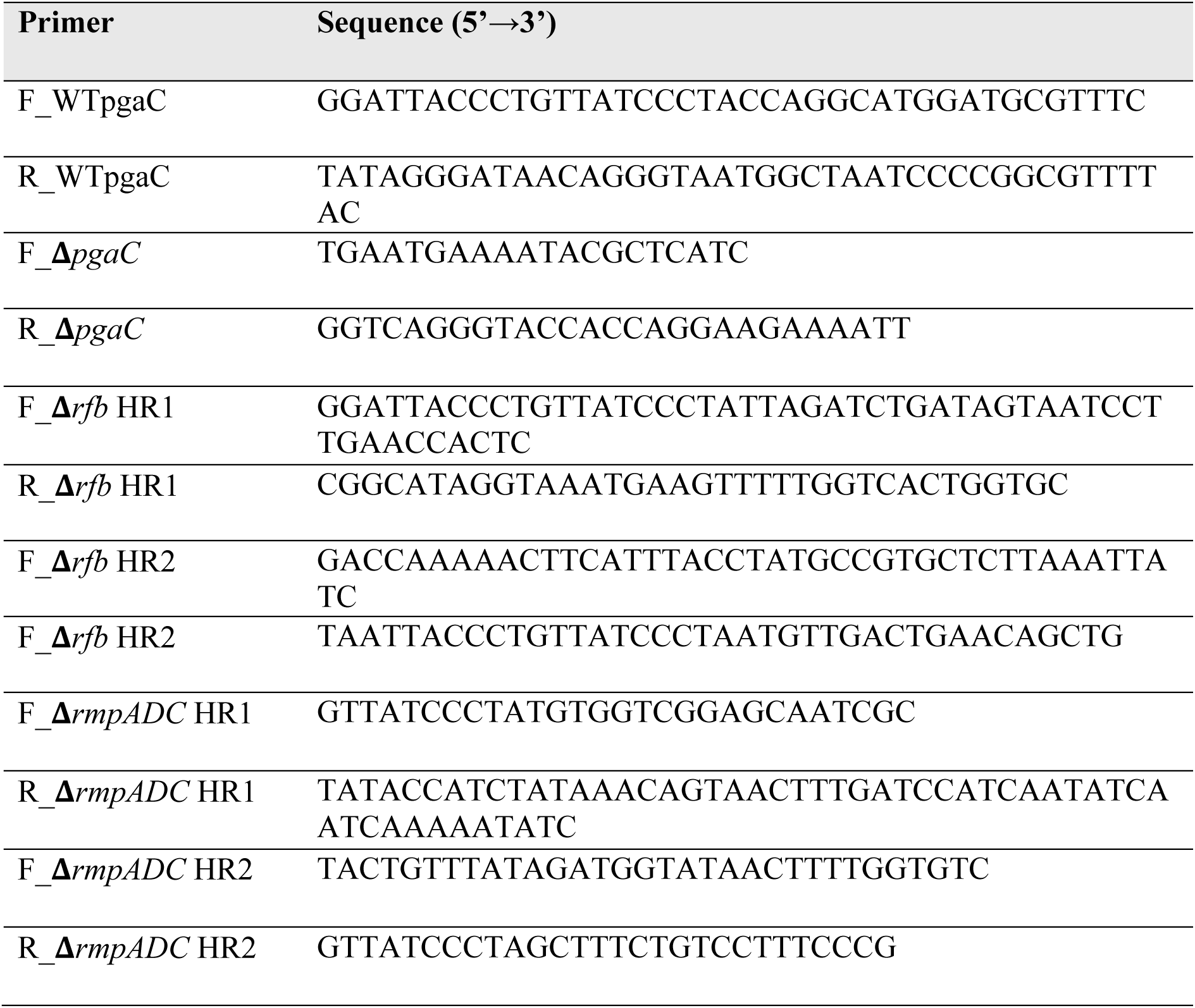
Primers used in this study.

**Table S4.**
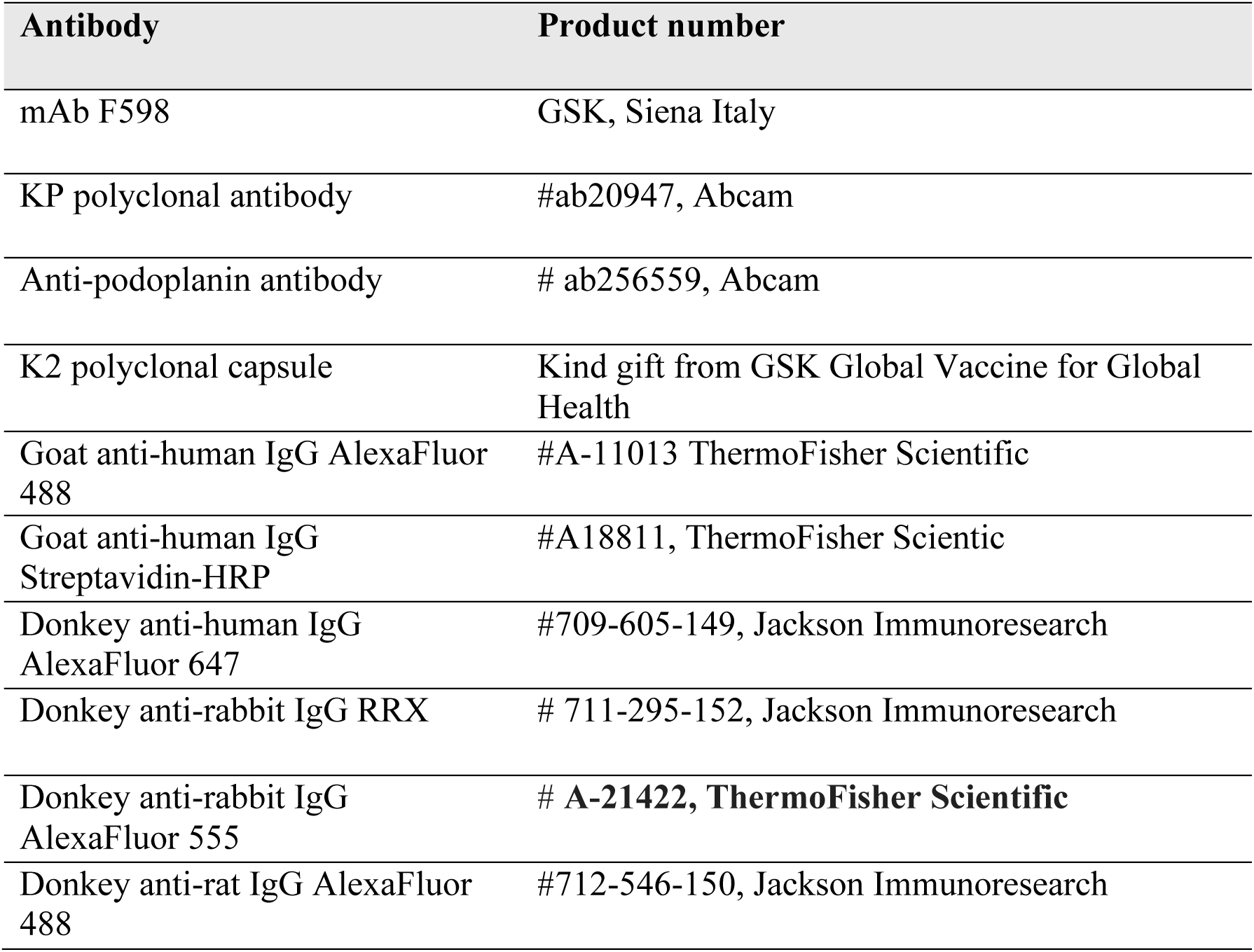
Antibodies used in this study.

## References

1. Little DJ, Li G, Ing C, DiFrancesco BR, Bamford NC, Robinson H, et al. Modification and periplasmic translocation of the biofilm exopolysaccharide poly-β-1, 6-N-acetyl-D-glucosamine. National Acad Sciences; 2014;111:11013–8.

2. Bentancor LV, O’Malley JM, Bozkurt-Guzel C, Pier GB, Maira-Litrán T. Poly-N-acetyl-β-(1-6)-glucosamine is a target for protective immunity against Acinetobacter baumannii infections. Am Soc Microbiol; 2012;80:651–6.

3. Gening ML, Maira-Litrán T, Kropec A, Skurnik D, Grout M, Tsvetkov YE, et al. Synthetic β-(1→ 6)-linked N-acetylated and nonacetylated oligoglucosamines used to produce conjugate vaccines for bacterial pathogens. Am Soc Microbiol; 2010;78:764–72.

4. David S, Wong JL, Sanchez-Garrido J, Kwong H-S, Low WW, Morecchiato F, et al. Widespread emergence of OmpK36 loop 3 insertions among multidrug-resistant clones of Klebsiella pneumoniae. Public Library of Science San Francisco, CA USA; 2022;18:e1010334.

5. Wong JL, Romano M, Kerry LE, Kwong H-S, Low W-W, Brett SJ, et al. OmpK36-mediated Carbapenem resistance attenuates ST258 Klebsiella pneumoniae in vivo. Nature Publishing Group UK London; 2019;10:3957.

6. Du Sert NP, Ahluwalia A, Alam S, Avey MT, Baker M, Browne WJ, et al. Reporting animal research: Explanation and elaboration for the ARRIVE guidelines 2.0. Public Library of Science; 2020;18:e3000411.

7. Choi AH, Slamti L, Avci FY, Pier GB, Maira-Litrán T. The pgaABCD locus of Acinetobacter baumannii encodes the production of poly-β-1-6-N-acetylglucosamine, which is critical for biofilm formation. Am Soc Microbiol; 2009;191:5953–63.

8. Low WW, Wong JL, Beltran LC, Seddon C, David S, Kwong H-S, et al. Mating pair stabilization mediates bacterial conjugation species specificity. Nature Publishing Group UK London; 2022;7:1016–27.

9. Berger CN, Crepin VF, Roumeliotis TI, Wright JC, Carson D, Pevsner-Fischer M, et al. Citrobacter rodentium subverts ATP flux and cholesterol homeostasis in intestinal epithelial cells in vivo. Elsevier; 2017;26:738–752. e6.

10. Podschun R, Ullmann U. Klebsiella spp. as nosocomial pathogens: epidemiology, taxonomy, typing methods, and pathogenicity factors. Am Soc Microbiol; 1998;11:589–603.

11. Delany I, Rappuoli R, Seib KL. Vaccines, reverse vaccinology, and bacterial pathogenesis. Cold Spring Harbor Laboratory Press; 2013;3:a012476.

12. Bleby J. Animals (Scientific Procedures) Act 1986. 1986;119:22.

13. Cywes-Bentley C, Skurnik D, Zaidi T, Roux D, DeOliveira RB, Garrett WS, et al. Antibody to a conserved antigenic target is protective against diverse prokaryotic and eukaryotic pathogens. National Acad Sciences; 2013;110:E2209–18.

14. Assoni L, Girardello R, Converso TR, Darrieux M. Current stage in the development of Klebsiella pneumoniae vaccines. Springer; 2021;10:2157–75.

15. Davenport ML, Sherrill TP, Blackwell TS, Edmonds MD. Perfusion and inflation of the mouse lung for tumor histology. 2020;e60605.

16. McKenney D, Pouliot KL, Wang Y, Murthy V, Ulrich M, Doring G, et al. Broadly protective vaccine for Staphylococcus aureus based on an in vivo-expressed antigen. American Association for the Advancement of Science; 1999;284:1523–7.

17. Pons S, Frapy E, Sereme Y, Gaultier C, Lebreton F, Kropec A, et al. A high-throughput sequencing approach identifies immunotherapeutic targets for bacterial meningitis in neonates. Elsevier; 2023;88.

18. Follador R, Heinz E, Wyres KL, Ellington MJ, Kowarik M, Holt KE, et al. The diversity of Klebsiella pneumoniae surface polysaccharides. Microbiology Society; 2016;2:e000073.

19. Wang X, Preston III JF, Romeo T. The pgaABCD locus of Escherichia coli promotes the synthesis of a polysaccharide adhesin required for biofilm formation. Am Soc Microbiol; 2004;186:2724–34.

20. Walker KA, Treat LP, Sepúlveda VE, Miller VL. The small protein RmpD drives hypermucoviscosity in Klebsiella pneumoniae. Am Soc Microbiol; 2020;11:10.1128/mbio.01750-20.

21. Skurnik D, Roux D, Pons S, Guillard T, Lu X, Cywes-Bentley C, et al. Extended-spectrum antibodies protective against carbapenemase-producing Enterobacteriaceae. Oxford University Press; 2016;71:927–35.

22. Maira-Litrán T, Kropec A, Abeygunawardana C, Joyce J, Mark III G, Goldmann DA, et al. Immunochemical properties of the staphylococcal poly-N-acetylglucosamine surface polysaccharide. Am Soc Microbiol; 2002;70:4433–40.

23. Opoku-Temeng C, Kobayashi SD, DeLeo FR. Klebsiella pneumoniae capsule polysaccharide as a target for therapeutics and vaccines. Elsevier; 2019;17:1360–6.

24. Martin MJ, Stribling W, Ong AC, Maybank R, Kwak YI, Rosado-Mendez JA, et al. A panel of diverse Klebsiella pneumoniae clinical isolates for research and development. Microbiology Society; 2023;9:000967.

25. Chen K-M, Chiang M-K, Wang M, Ho H-C, Lu M-C, Lai Y-C. The role of pgaC in Klebsiella pneumoniae virulence and biofilm formation. Elsevier; 2014;77:89–99.

26. Singh S, Wilksch JJ, Dunstan RA, Mularski A, Wang N, Hocking D, et al. LPS O antigen plays a key role in Klebsiella pneumoniae capsule retention. Am Soc Microbiol; 2022;10:e01517–21.

27. Wong JL, David S, Sanchez-Garrido J, Woo JZ, Low WW, Morecchiato F, et al. Recurrent emergence of Klebsiella pneumoniae carbapenem resistance mediated by an inhibitory ompK36 mRNA secondary structure. National Acad Sciences; 2022;119:e2203593119.

28. David S, Reuter S, Harris SR, Glasner C, Feltwell T, Argimon S, et al. Epidemic of carbapenem-resistant Klebsiella pneumoniae in Europe is driven by nosocomial spread. Nature Publishing Group UK London; 2019;4:1919–29.

29. Walker KA, Miner TA, Palacios M, Trzilova D, Frederick DR, Broberg CA, et al. A Klebsiella pneumoniae regulatory mutant has reduced capsule expression but retains hypermucoviscosity. Am Soc Microbiol; 2019;10:10.1128/mbio.00089-19.

30. Ernst CM, Braxton JR, Rodriguez-Osorio CA, Zagieboylo AP, Li L, Pironti A, et al. Adaptive evolution of virulence and persistence in carbapenem-resistant Klebsiella pneumoniae. Nature Publishing Group US New York; 2020;26:705–11.

31. Hu F, Pan Y, Li H, Han R, Liu X, Ma R, et al. Carbapenem-resistant Klebsiella pneumoniae capsular types, antibiotic resistance and virulence factors in China: a longitudinal, multi-centre study. Nature Publishing Group UK London; 2024;9:814–29.

32. Dai P, Hu D. The making of hypervirulent Klebsiella pneumoniae. Wiley Online Library; 2022;36:e24743.

33. Itoh Y, Rice JD, Goller C, Pannuri A, Taylor J, Meisner J, et al. Roles of pgaABCD genes in synthesis, modification, and export of the Escherichia coli biofilm adhesin poly-β-1, 6-N-acetyl-D-glucosamine. Am Soc Microbiol; 2008;190:3670–80.

34. Hopkins EG, Roumeliotis TI, Mullineaux-Sanders C, Choudhary JS, Frankel G. Intestinal epithelial cells and the microbiome undergo swift reprogramming at the inception of colonic Citrobacter rodentium infection. Am Soc Microbiol; 2019;10:10.1128/mbio.00062-19.

35. Bain W, Ahn B, Peñaloza HF, McElheny CL, Tolman N, van der Geest R, et al. In Vivo Evolution of a Klebsiella pneumoniae Capsule Defect With wcaJ Mutation Promotes Complement-Mediated Opsonophagocytosis During Recurrent Infection. Oxford University Press; 2024;jiae003.

36. Steiner S, Lori C, Boehm A, Jenal U. Allosteric activation of exopolysaccharide synthesis through cyclic di-GMP-stimulated protein–protein interaction. John Wiley & Sons, Ltd Chichester, UK; 2013;32:354–68.

37. Bengoechea JA, Sa Pessoa J. Klebsiella pneumoniae infection biology: living to counteract host defences. Oxford University Press; 2019;43:123–44.

38. Walter J, Haller S, Quinten C, Kärki T, Zacher B, Eckmanns T, et al. Healthcare-associated pneumonia in acute care hospitals in European Union/European Economic Area countries: an analysis of data from a point prevalence survey, 2011 to 2012. European Centre for Disease Prevention and Control; 2018;23:1700843.

39. WHO Bacterial Priority Pathogens List, 2024.

40. Yu W-L, Ko W-C, Cheng K-C, Lee C-C, Lai C-C, Chuang Y-C. Comparison of prevalence of virulence factors for Klebsiella pneumoniae liver abscesses between isolates with capsular K1/K2 and non-K1/K2 serotypes. Elsevier; 2008;62:1–6.

41. Tan Z, Yang W, O’Brien NA, Pan X, Ramadan S, Marsh T, et al. A comprehensive synthetic library of poly-N-acetyl glucosamines enabled vaccine against lethal challenges of Staphylococcus aureus. Nature Publishing Group UK London; 2024;15:3420.

